# Weakly supervised identification of microscopic human breast cancer-related optical signatures from normal-appearing breast tissue

**DOI:** 10.1101/2022.05.24.493356

**Authors:** Jindou Shi, Haohua Tu, Jaena Park, Marina Marjanovic, Anna M. Higham, Natasha N. Luckey, Kimberly A. Cradock, Z. George Liu, Stephen A. Boppart

## Abstract

With the latest advancements in optical bioimaging, rich structural and functional information has been generated from biological samples, which calls for capable computational tools to identify patterns and uncover relationships between optical characteristics and various biomedical conditions. Constrained by the existing knowledge of the novel signals obtained by those bioimaging techniques, precise and accurate ground truth annotations can be difficult to obtain. Here we present a weakly supervised Deep Learning framework for optical signature discovery based on inexact and incomplete supervision. The framework consists of a Multiple Instance Learning-based classifier for the identification of regions of interest in coarsely labeled images, and model interpretation techniques for optical signature discovery. We applied this framework to investigate human breast cancer-related optical signatures based on virtual histopathology enabled by simultaneous label-free autofluorescence multiharmonic microscopy (SLAM), with the goal to explore unconventional cancer-related optical signatures from normal-appearing breast tissues. The framework has achieved an average area under the curve (AUC) of 0.975 on the cancer diagnosis task. In addition to well-known cancer biomarkers, non-obvious cancer-related patterns were revealed by the framework, including NAD(P)H-rich extracellular vesicles observed in normal-appearing breast cancer tissue, which facilitate new insights into the tumor microenvironment and field cancerization. This framework can be further extended to diverse imaging modalities and optical signature discovery tasks.

## 1. Introduction

Advancements in optical bioimaging have revealed rich structural and functional information from biological samples based on intrinsic structural, molecular, and metabolic contrasts [1–4]. Without requiring tissue fixation, sectioning, and staining, label-free virtual histopathology technology fulfilled by multimodal optical bioimaging techniques enables the observation of the unperturbed biochemical microenvironment [5, 6]. Based on the collected morphological and molecular information, unconventional cancer-related optical signatures can be revealed, which are mainly undetectable using traditional histopathological techniques. However, the rich information in multimodal optical bioimages may easily exceed the capabilities of human visual inspection. Extracting information and detecting patterns using computational tools could therefore enhance perceptions and inspire new scientific discoveries.

Deep Learning (DL), which excels at extracting patterns and insights from large volume of data, has achieved substantial success in biomedical image analysis [7–9]. However, generating accurate and fine-grained manual annotations for DL model training can be time-consuming, costly, and sometimes unfeasible [10]. To alleviate the dependency on the strong supervision, numerous weakly supervised learning approaches have been proposed to exploit inexact and incomplete supervision [11–14]. Among those approaches, Multiple Instance Learning (MIL), which leverages inexact supervision, has gained popularity in computational pathology where the generation of fine-grained annotations are often costly and time-consuming [15–18]. Instead, coarse annotations (e.g., labels for the whole slides images) are available more readily. It was reported that MIL-based decision support systems have achieved clinical-grade performance for cancer diagnosis and identified cancer-related regions when being trained on large-size histology datasets without cellular-level annotations [19].

In this study, we leverage MIL for human breast cancer-related optical signature identification based on label-free virtual histopathology, where only ambiguous whole-image-level annotations are available. The virtual histopathology slides were generated using simultaneous label-free autofluorescence-multiharmonic (SLAM) microscopy which enables simultaneous and efficient acquisition of two- and three-photon-excited autofluorescence (2PF and 3PF, respectively) as well as second and third harmonic generation (SHG and THG, respectively) with strict spatial and temporal co-registration (Fig. 1) [6, 20]. In addition to structural information, SLAM provides the molecular information of a sample in its native state, demonstrating its great potential for the exploration of novel molecular biomarkers at early stages of cancer, as well as for new fundamental investigations into carcinogenesis [20, 21].

**Fig. 1.**
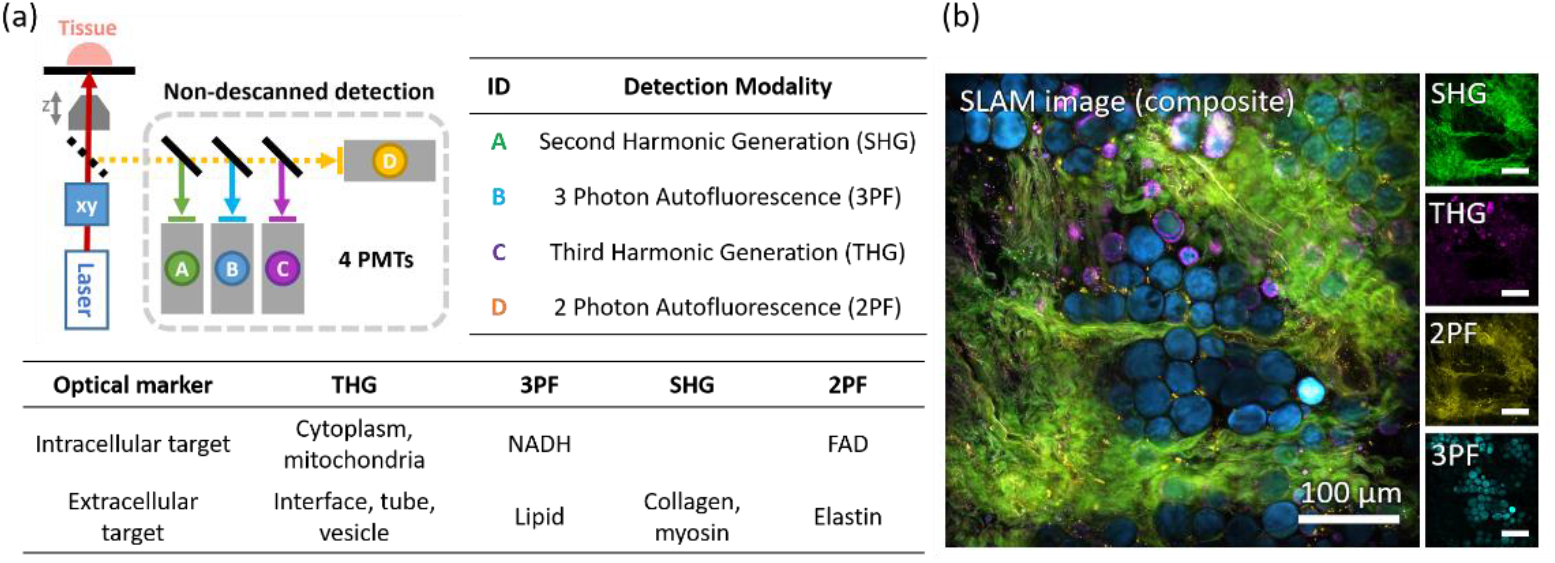
Description of the SLAM imaging system and a sample SLAM virtual histopathology slide image. (a) Overview of SLAM imaging system, with the schematic of SLAM microscope (upper left), detection modalities (upper right table), and the targeted intracellular and extracellular optical markers (bottom table). (b) SLAM virtual histopathology slide image collected in the tumor adjacent region of a breast tissue from a cancer human subject.

Unlike many DL-based cancer diagnosis applications which focus primarily on tissue regions with well-established cancer-associated morphologies, this study aims to explore unconventional structural and functional optical signatures for which the correlations with breast cancer might not be fully investigated. In addition to primary tumors, the SLAM virtual histopathology slide images (“slides” for brevity) of tumor microenvironment (TME) and peritumoral regions were generated to reveal potential evidence of field cancerization [22, 23]. Studies have shown various interactions between tumor and TME linked with tumor angiogenesis and peripheral immune tolerance, which may affect the growth and evolution of cancerous cells [24]. The replacement of large areas of normal cell population by cancer-primed cell population is often referred to field cancerization that may not involve clinically detectable morphological changes [25].

However, constrained by the current knowledge of the unconventional functional and structural information generated by SLAM, the annotations for the whole images can be ambiguous. Namely, tissue images from tumor surrounding regions without conventional cancer biomarkers can hardly be assigned certain labels (cancer or normal). Additionally, the cancer-related optical signatures from uncertain regions (e.g., tumor surrounding areas) may differ from those inside the well-known cancer areas (i.e., primary tumor). This leads to a unique learning task where DL models need to be trained with inexact and incomplete supervision at the same time (Fig. S1 in Supplement 1). Therefore, a DL framework that exploits weak supervision is needed for discovery of cancer-related patterns in both the primary tumor regions and tumor surrounding regions where potential optical markers for field cancerization and carcinogenesis may exist. In this study, the cancer-related patterns are defined as optical characteristics exhibited only in tissues from cancer subjects while being unobservable in tissues from normal subjects.

Here, we propose Mix-and-Match Multiple Instance Learning (MM-MIL) as a weakly supervised DL framework for the discovery of human breast cancer-related optical signatures. By modifying the bagging policy in conventional MIL, MM-MIL is designed to learn from whole-image-level ambiguous annotations. Through iterative selection and learning of discriminative instances from positive and negative bags, MM-MIL could make whole-slide level cancer diagnosis, as well as inform cancer-related regions in predicted positive slides. In addition to well-established cancer biomarkers, MM-MIL revealed non-obvious cancer-related signatures that appear predominantly in tumor surrounding areas while being unobservable in breast tissues from normal subjects. Those optical signatures may inspire the discovery of new cancer markers inside and outside of the tumor microenvironment and promote research on carcinogenesis, tumor progression, and field cancerization.

## 2. Materials and methods

### 2.1 SLAM virtual histopathology dataset generation

SLAM virtual histopathology slides were collected from fresh human breast tissue specimens using a custom-built benchtop SLAM microscope [20], and under a protocol approved by the Institutional Review Boards at Carle Foundation Hospital and the University of Illinois at Urbana-Champaign. Tissue samples from 48 human subjects was used in this study, among which 36 subjects were diagnosed with ductal carcinoma in situ (DCIS) or invasive breast cancer (invasive ductal carcinoma, invasive lobular carcinoma, lobular carcinoma in situ, invasive papillary carcinoma) by board-certified pathologists. Normal tissue samples were collected from 12 subjects undergoing breast reduction surgery, with no reported history of cancer. Information on subject demographics can be found in Table S1 in Supplement 1.

We generated 724 SLAM slides, where each slide is a mosaic image with a median size of 1500 × 1500 pixels (750 μm × 750 μm). To investigate the potential optical signatures for field cancerization, both tumor and tumor adjacent regions in cancer tissue samples were imaged. For images collected in tumor adjacent regions, the distances between the imaging site and the tumor boundary (annotated by pathologists) ranged from several millimeters to several centimeters. Slide-level annotations were generated by human experts based on pathological reports, optical characteristics in SLAM, and the corresponding hematoxylin-eosin (H&E) histological slides (Fig. S2). Slides that exhibit well-known cancer biomarkers (e.g., tumor cells, tumor-associated collagen fibers) were annotated as “Cancer”. “Uncertain” labels were assigned to slides that were from cancer subjects but do not show well-known cancer signatures (mainly outside the primary tumor). Slides from normal subjects were labeled as “Normal”. The dataset was randomly split into a training set (about 60% of slides), a validation set (about 10% of slides), and a test set (about 30% of slides) using subject-level partition such that slides from one subject can only exist in one of these sets. A description of slides in each set is provided in Table 1.

**Table 1.** Description of the SLAM virtual histopathology dataset.

| Dataset | Total subjects | Cancer subjects | SLAM slides | “Cancer” slides | “Uncertain” slides | “Normal” slides | Size (Mpixels) |
| --- | --- | --- | --- | --- | --- | --- | --- |
| Training | 23 | 17 | 428 | 59 | 237 | 132 | 1,077 |
| Validation | 7 | 5 | 92 | 7 | 61 | 24 | 203 |
| Test | 18 | 14 | 204 | 23 | 137 | 44 | 484 |
| Total | 48 | 36 | 724 | 89 | 435 | 200 | 1,764 |

### 2.2 SLAM slide preprocessing

Each channel of SLAM slides was normalized using Z-score standardization. In addition, slides were augmented by flips and rotations: horizontal flip, vertical flip, 90-degree rotation, 180-degree rotation, 270-degree rotation. Slides were then cropped into tile images (“tiles” for simplicity), which share the same label as the slides. To investigate cancer-related patterns at different scales, three tile sizes were used to simulate high-, medium-, and low-magnification levels. For the high-magnification level, slides were cropped to 256 × 256-pixel tiles (128 μm × 128 μm) with no overlap between tiles. Medium- and low-magnification levels tiles were generated by tiling slides into 512 × 512-pixel images (256 μm × 256 μm), and 1024 × 1024-pixel images (512 μm × 512 μm), respectively, and downscaling to 256 × 256-pixel tiles was performed using nearest-neighbor interpolation. These tiles cover a larger FOV, but have lower resolution compared to high-magnification tiles. Overlap ratios of 50% and 80% were used for medium- and low-magnification levels, respectively. The numbers of tiles at different magnification levels are reported in Table S2 in Supplement 1.

### 2.3 MM-MIL framework

To identify and localize cancer-related patterns in coarsely labeled SLAM slides, the cancer signature discovery task was formulated a MIL problem, where SLAM tiles were treated as MIL instances. To enable the MIL model to extract cancer-related patterns from both “Cancer” and “Uncertain” slides, a custom-designed bagging policy (i.e., Mix-and-Match) was developed to ensure the explicitness of bag-level labels (Fig. 2). Instead of treating each SLAM slide as a MIL bag, slides were mixed and grouped together to form bags in MIL. For each set (i.e., training, validation, or test set), a positive bag was defined as a group of “Cancer” slides and “Uncertain” slides. Since at least one tile in the positive bag has cancer signatures, explicit labels (“Positive”) could be assigned to the bag based on the standard assumption of MIL [26]. Considering that “Normal” slides came from normal subjects with no history of cancer, each of them can form an individual bag with a certain “Negative” label. To maximize the number of bags, every positive bag contained only one “Cancer” slide. While “Uncertain” slides were randomly distributed to all positive bags. Notably, only bag-level labels were used for model training, while slide-level labels were hidden to encourage models to look for cancer-related optical signatures in both “Cancer” and “Uncertain” slides.

**Fig. 2.**
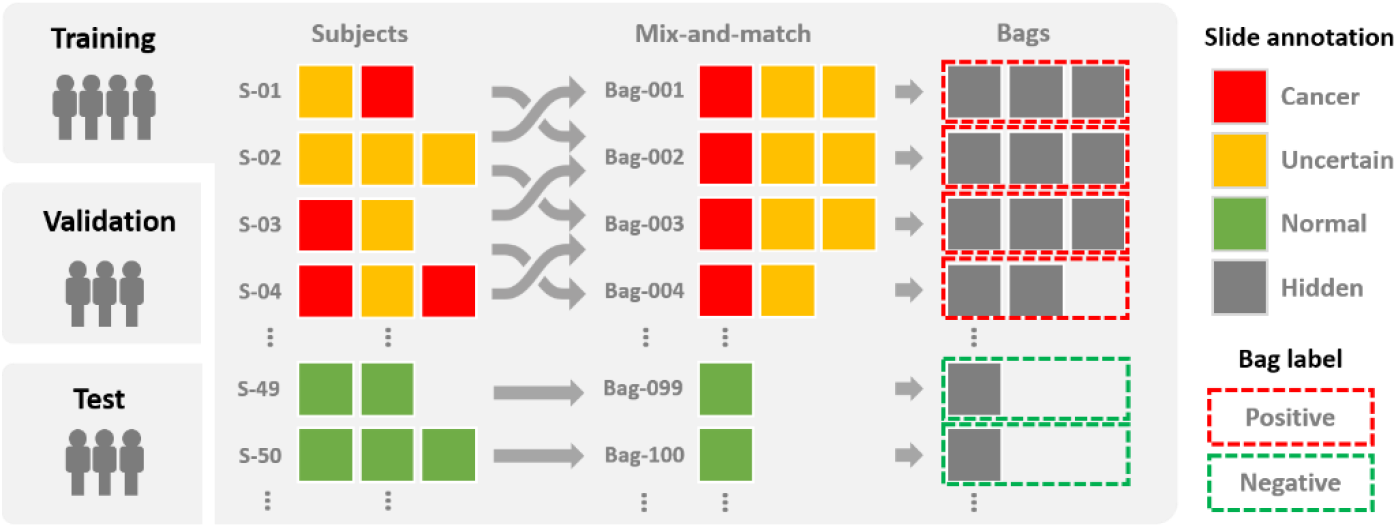
The Mix-and-Match bagging policy in MM-MIL. “Cancer” slides and “Uncertain” slides were randomly grouped together to form positive bags, whereas “Normal” slides were treated as negative bags. During model training, all slide-level labels were hidden, and only bag-level labels (“Positive” or “Negative”) were used.

Based on the MIL bags generated by the Mix-and-Match bagging policy, DL-based models were trained to make cancer diagnosis on the bag level (i.e., global prediction) as well as on the instance level (i.e., local prediction). This was achieved by training an instance-level classifier and aggregating the instance-level predictions to generate bag-level classification results (Fig. 3). Inspired by [19, 27], the instance-level classifier (*f_ins_*) was trained in an iterative manner using the EM algorithm. Each training epoch started with an inference step (the E-step in EM), where the model generated prediction scores (i.e., cancer probability) for all the tiles in every training bag. The training sample pool *T_train_* was then constructed by selecting the top *K* instances with highest prediction scores in each bag:

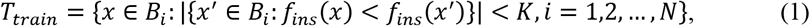

where *x* represents an instance, *B_i_* is the *i*-th bag among the total *N* bags, and *K* is a user-defined parameter. The top *K* instances shared the labels with their bags. The instance classifier was then trained on the training sample pool *T_train_* for one epoch (i.e., the M-step in EM). The training procedure of the instance-level classifier is further illustrated in Supplement 1. In this study, we use the ResNet34 as the backbone of the instance-level classifier [28]. The model was initialized using weights pretrained on ImageNet except for the first convolution layer and the last fully connected layer, of which the input and output dimensions were customized for the task, were randomly initialized.

**Fig. 3.**
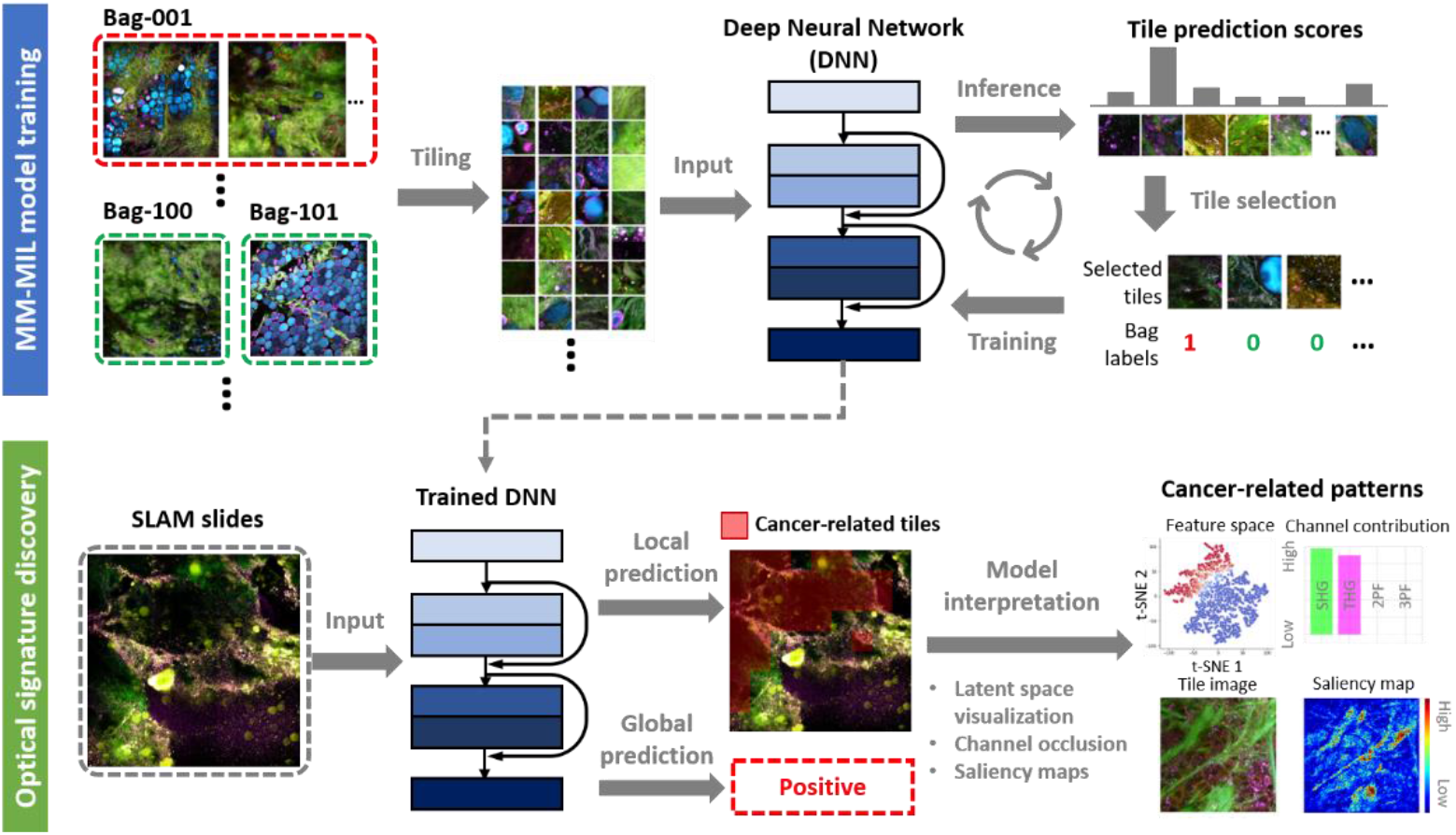
Description of the MM-MIL framework. During model training, a DNN-based tile-level classifier is optimized through iterative selection and learning of discriminative tiles from Mix- and-Match bags. Once the classifier is trained, local and global cancer predictions can be generated. Model interpretation methods are utilized to inform cancer-related patterns learned by the model.

Bag-level predictions were obtained by aggregating instance-level prediction scores. Constrained by the size of the dataset used in this study, we chose a nonparametric method *f_bag_* (i.e., max-pooling operation) for bag-level aggregation. That is, if at least one instance in a particular bag *B_i_* is positive (probability > 0.5), the whole bag is predicted as positive:

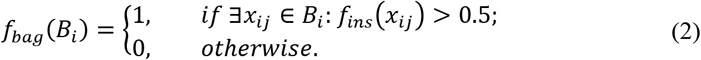

Compared to trainable bag aggregation methods [29, 30], the max-pooling operation is more sensitive to spurious positive instances, thus imposes higher requirements on the reliability of instance-level classifiers.

### 2.3 Training strategy and loss function

Each MM-MIL instance classifier was trained for at least 50 epochs and at most 200 epochs. Model performance was evaluated on the validation set, where the classification accuracy was recorded. Early stopping would be triggered when the validation accuracy did not increase for 30 consecutive epochs. The model with highest validation accuracy was saved for final performance evaluation on the test set. The objective function of the instance-level classifier is the cross-entropy loss:

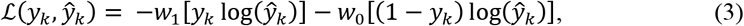

where *y_k_*, *ŷ_k_* are the bag-level ground truth and prediction score respectively, and *w_0_*, *w_1_* are negative and positive class weights, which are set to deal with the unbalanced class frequency within the training sample pool. The optimization of model parameters was achieved by stochastic gradient descent using the Adam optimizer, with a learning rate of 1.5 × 10^-4^, and a weight decay of 1 × 10^-5^. A detailed description of hyperparameters used in the experiments can be found in Table S3 in Supplement 1.

### 2.3 Model evaluation

The global prediction generated by MM-MIL were evaluated as a binary classification task. The average performance of five MM-MIL models trained with the same configuration was reported. For each magnification level (i.e., high, medium, and low), the receiver operating characteristic (ROC) curves and the associated area under the curve (AUC) were calculated. Due to the absence of certain ground truth labels for “Uncertain” slides, two approaches were adopted to evaluate the global prediction performance. For the first approach, MM-MIL models were evaluated on the Mix-and-Match bags in the test set, each of which has a certain “Positive” or “Negative” label. In the second approach, the model performance is assessed on the test slides with certain labels (i.e., “Cancer” and “Normal”).

To quantitatively evaluate the local (tile-level) predictions, known tumor areas in “Cancer” slides were annotated by human experts based on the appearance of known cancer signatures. Considering that the areas outside human-annotated tumor regions may also exhibit non-obvious cancer-related patterns, commonly used evaluation metrics (e.g., Intersection over Union (IoU)) may not be suitable in this study. Instead, the coverage ratio *R_coverage_* was used to measure the coverage of the model’s cancer predictions on known tumor regions. The known tumor regions in “Cancer” slides in the test set were annotated based on the well-known cancer signatures. The coverage ratio was calculated by *R_coverage_* = *m_pp_*/*m_TP_*, where *m_TP_* is the number of tumor tiles, of which at least 50% of the tile area is annotated as tumor region, whereas *m_TP_* is the number of predicted positive tiles among the *m_TP_* tumor tiles. In addition, the positive tile ratios in each group of slides (i.e., “Cancer”, “Uncertain”, and “Normal”) were calculated by *R_positive_* = *m_positive_*/*m_total_* where the total number of tiles and the number of positive tiles in a particular group are denoted as *m_total_*, and *m_positive_* respectively.

### 2.4 Model interpretation

To gain insights into model predictions, three model interpretation methods were leveraged for the discovery of cancer-related optical signatures. Firstly, the latent feature space of the ResNet34 model was visualized in two dimensions using t-distributed stochastic neighbor embedding (t-SNE) [31]. Tiles (converted to RGB images) corresponding to the points in the t-SNE plot were sampled and visualized for model introspection. Secondly, SLAM channels of predicted positive tiles was occluded to investigate the importance of each channel for the cancer classification task. The decrease in prediction score can be seen as an indicator of channel importance regarding to the prediction task. Lastly, saliency map techniques, Integrated Gradients (IG) [32] and SmoothGrad (SG) [33], were utilized to inform the salient structures on the pixel-level. The implementation details of model interpretation were reported in Supplement 1.

### 2.5 Implementation details

All data processing and model development were conducted on a workstation computer, equipped with an Intel Xeon W-2195 central processing unit (CPU), four Nvidia RTX 8000 graphics processing units (GPUs), and 256 gigabytes of memory. The workstation operates on the Ubuntu system (version 18.04). During the model optimization, each batch (i.e., 600 tiles per batch) required 0.365 s for the inference step, and 0.694 s for the training step with four GPUs. At the prediction time, MM-MIL generates diagnoses at an average speed of 23.39 slides/s in high-magnification mode, and 26.17 slides/s in low-magnification mode on 2000×2000-pixel SLAM slides. The source code for MM-MIL is available in a public repository on GitHub, https://github.com/Biophotonics-COMI/MM-MIL.

## 3. Results

### 3.1 Global prediction evaluation

The global prediction evaluation results are shown in Fig. 4(a)(b) and Table 2. MM-MIL achieved average AUCs of 0.975, 0.971, and 0.969 when evaluated on the Mix-and-Match bags under high-, medium-, and low-magnification settings respectively [Fig. 4(a)]. In addition, when evaluated on the test slides with certain labels (“Cancer” or “Normal”), an average AUC of 0.939 was achieved for the high-magnification level. While for medium- and low-magnification settings, average AUCs of 0.810 and 0.816 were achieved respectively [Fig. 4(b)]. The differences in AUCs between bag- and slide-level predictions indicate that the “Uncertain” slides in positive bags have considerable contribution to cancer prediction. For both bag-level and slide-level global prediction tasks, we observed that MM-MIL models trained on the high magnification level achieved the best performance.

**Fig. 4.**
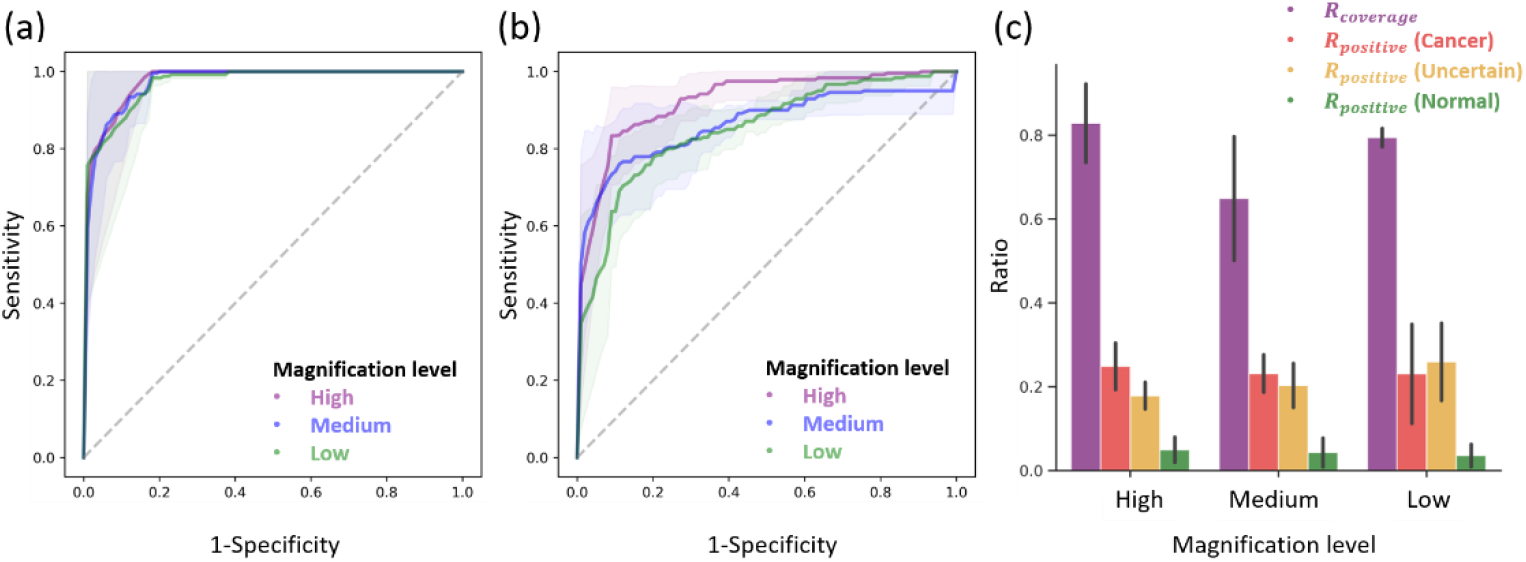
Global and local prediction evaluation results. ROC curves for MM-MIL models evaluated on the Mix-and-Match bags (a) and the slides with certain labels (b). The solid lines indicate the mean curve over five models trained under the same configuration, and the shaded regions indicate ± standard deviation. (c) Bar chart of local prediction evaluation metrics. The error bars indicate ± standard deviation.

**Table 2.** Global and local evaluation metrics (mean ± standard deviation) for MM-MIL models trained on high-, medium-, and low-magnification levels

| Magnification level | Global prediction evaluation |  | Local prediction evaluation |  |  |  |
| --- | --- | --- | --- | --- | --- | --- |
| | Bag-level AUC | Slide-level AUC | $R_{coverage}$ | $R_{positive}(\text{cancer})$ | $R_{positive}(\text{uncertain})$ | $R_{positive}(\text{normal})$ |
| High | **0.975±0.033** | **0.924±0.040** | **0.823±0.094** | 0.249±0.062 | 0.178±0.036 | 0.049±0.033 |
| Medium | 0.971±0.025 | 0.862±0.046 | 0.629±0.148 | 0.231±0.050 | 0.203±0.059 | 0.043±0.039 |
| Low | 0.969±0.041 | 0.850±0.044 | 0.794±0.022 | 0.231±0.132 | 0.259±0.103 | 0.036±0.030 |

In addition, experiments were conducted to compare the performance of model trained with the Mix-and-Match bagging policy and conventional bagging strategies (i.e., slide-level and subject-level). For the slide-level bagging policy, each slide was treated as a MIL bag. Slides from cancer subjects were labeled as positive. While for subject-level bagging policy, slides from individual subjects were grouped into a MIL bag with labels being determined by the subject information (positive for cancer subjects, and negative for normal subjects). Due to the fact that SLAM slides from cancer subjects might came from tumor surrounding regions, where evidence of cancer may not present, the ground truth labels for slide-level and subject-level bags can be unreliable. When being tested on the same held-out test set with certain slide labels, the MIL models achieved an average AUC of 0.939, 0.808, and 0.781 with Mix-and-Match, slide-level, and subject-level bagging strategies respectively (Fig. S3). In addition, the large variance in AUCs was observed for slide-level and subject-level bagging strategies, with the standard derivation to be 0.020, 0.211, and 0.257 for the three bagging strategies, respectively.

### 3.2 Local prediction evaluation

To evaluate the local (tile-level) predictions, cancer-related regions predicted by MM-MIL were compared with the known tumor areas with well-known cancer signatures. The resulting *R_coverage_* ratio was reported in Fig. 4(c) and Table 2. For models trained on the high-magnification mode, 82.3% of annotated-positive tiles were predicted as positive on average, while the models achieved average *R_coverage_* ratios of 62.9% and 79.4% for medium- and low-magnification modes respectively. *R_positive_* was calculated for “Cancer”, “Uncertain”, and “Normal” slides individually. For “Normal” slides from healthy subjects, whose tissue samples were not expected to show evidence of cancer, the average *R_positive_* ratios are 4.9%, 4.3%, and 3.6% under the high-, medium-, and low-magnification settings, respectively. Interestingly, a good number of tile images in “Uncertain” slides were predicted as positive, with average *R_positive_* ratios to be 17.8%, 20.3%, and 25.9% for the corresponding magnification levels, indicating that “Uncertain” slides may show evidence of cancer according to the MM-MIL models. However, from the perspective of human experts, the positive tiles in “Uncertain” slides do not exhibit well-known cancer-associated patterns, thus requiring further analysis and interpretation.

Overall, MM-MIL models trained on the high-magnification level achieved the highest AUC scores for the global prediction tasks, as well as highest *R_coverage_* for the local prediction tasks. In the following sections, we will focus on the high magnification level which has the most detailed local information about the tissue sample.

**Fig. 5.**
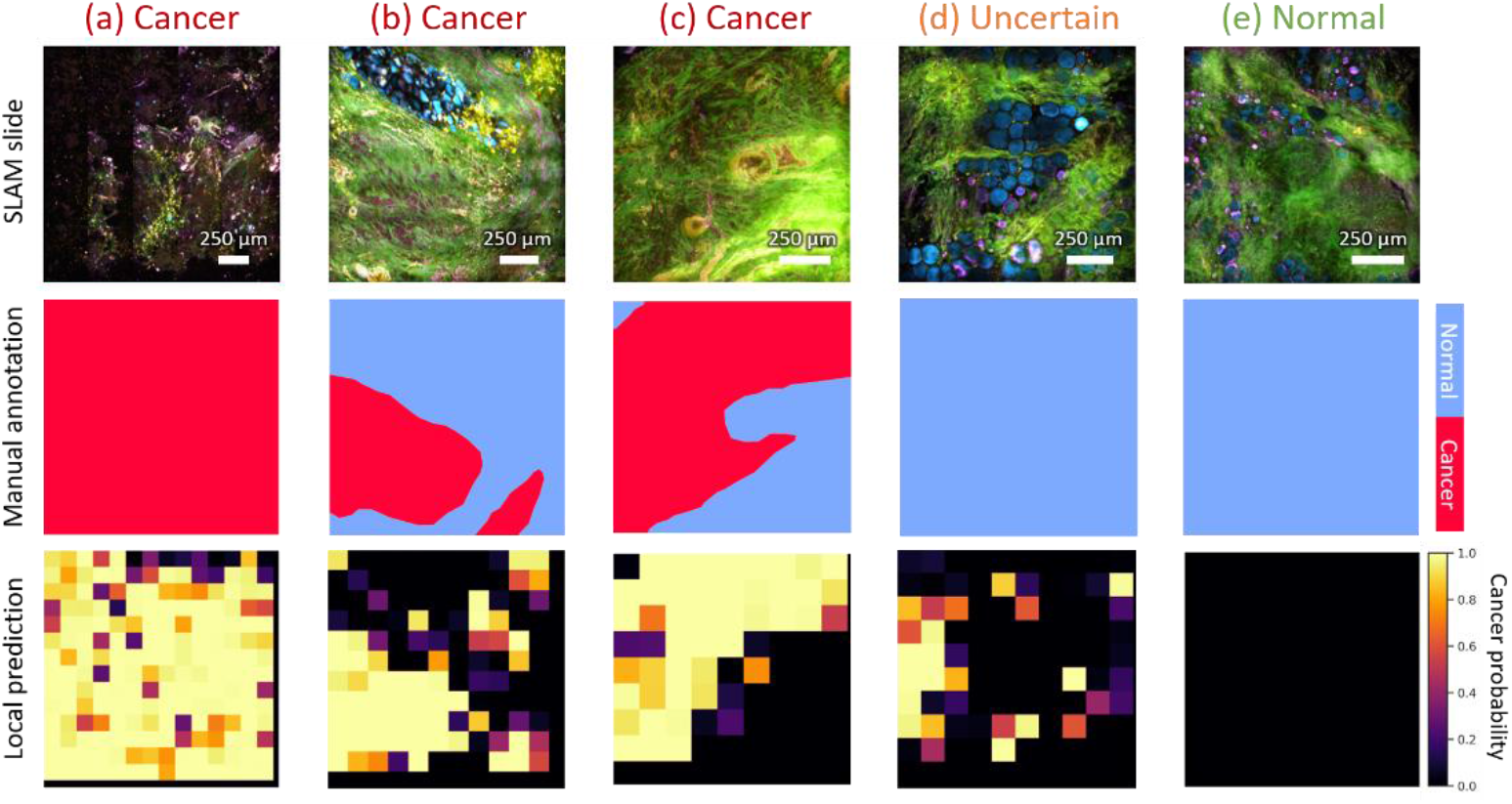
Cancer localization predictions of SLAM slides from “Cancer” group (a)-(c), “Uncertain” group (d), and “Normal” group (e) produced by a MM-MIL model trained on the high-magnification setting. The cancer regions were annotated by human experts based on well-known cancer biomarkers (second row). The cancer probabilities of each tile in the slide are visualized as heatmaps in the third row.

### 3.3 Visualization of latent feature space for model introspection

To gain insight into model predictions, the latent feature representations of tiles were visualized in two dimensions using t-SNE. Tiles with similar feature representations according to the model were grouped close to each other. Fig. 6 shows the t-SNE visualization of tiles from all test slides, and the t-SNE visualization of predicted positive tiles is shown in Fig. 7. As shown in Fig. 6(b), the majority of tiles from “Normal” slides have low cancer probability according to MM-MIL, whereas a considerable number of tiles from “Uncertain” slides were predicted positive (cancer probability > 0.5), as shown in Fig. 6(c). Noticeably, a large proportion (around 70%) of positive tiles from “Uncertain” slides formed separate clusters with positive tiles from “Cancer’ slides in the latent feature space, indicating that “Uncertain” slides may exhibit cancer-related patterns that were undetected in “Cancer” slides [Fig. 6(c)(d)]. Such cancer-related patterns were not observed in “Normal” slides [Fig. 6(b)], which implies that, based on SLAM microscopy, unique cancer-related patterns may be revealed from tumor-adjacent and peri-tumoral regions.

**Fig. 6.**
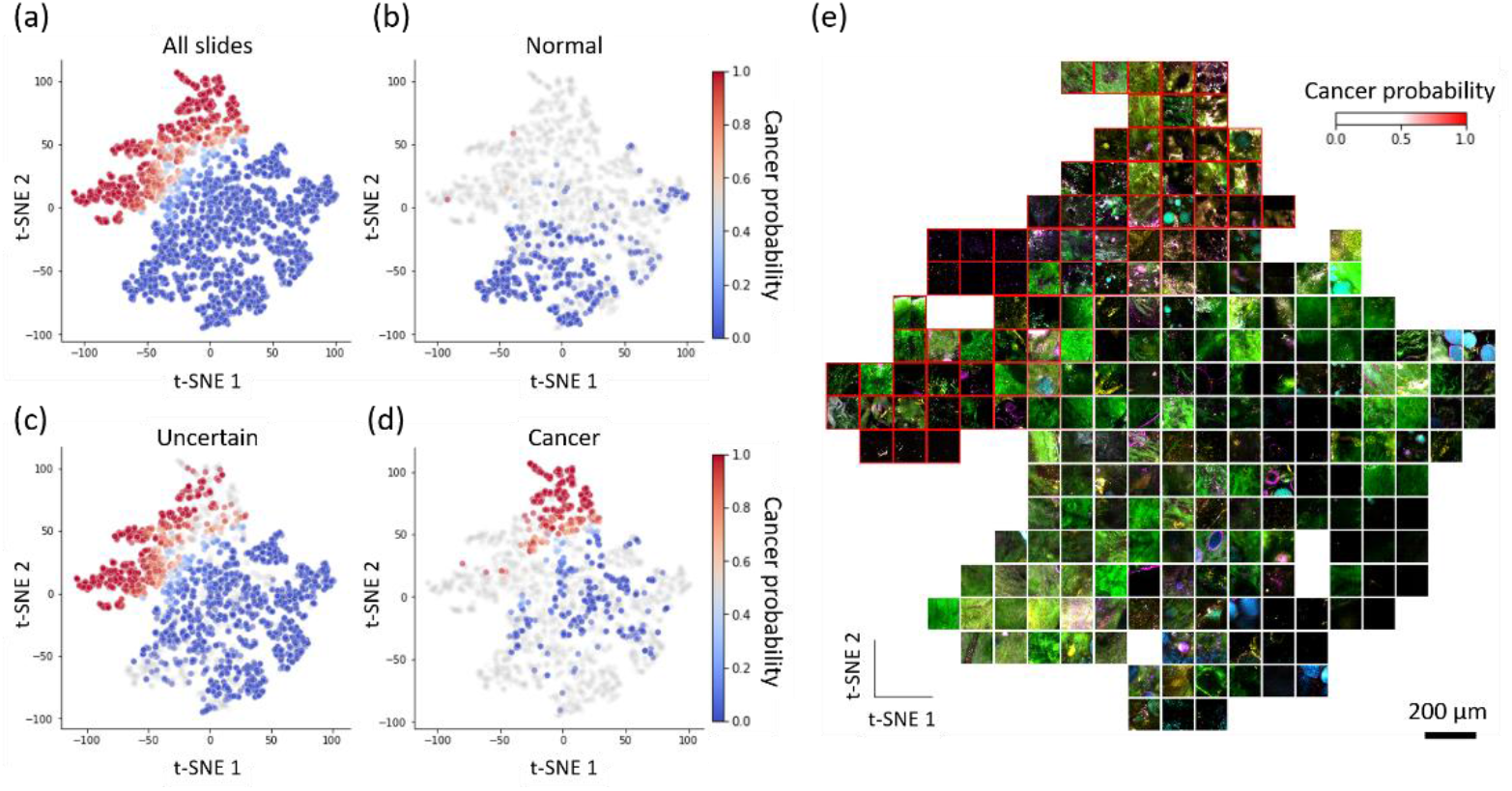
t-SNE visualization of the latent feature space of a trained MM-MIL model. (a) t-SNE plot of tiles from all slides in the test set. Each dot represents a SLAM tile image. Dots are colored based on the cancer probabilities predicted by the MM-MIL model. Tiles from “Normal”, “Uncertain”, and “Cancer” slides are highlighted in (b-d) separately. (e) SLAM tiles (composite images) corresponding to the dots in the t-SNE plot were sampled and visualized in the same locations.

**Fig. 7.**
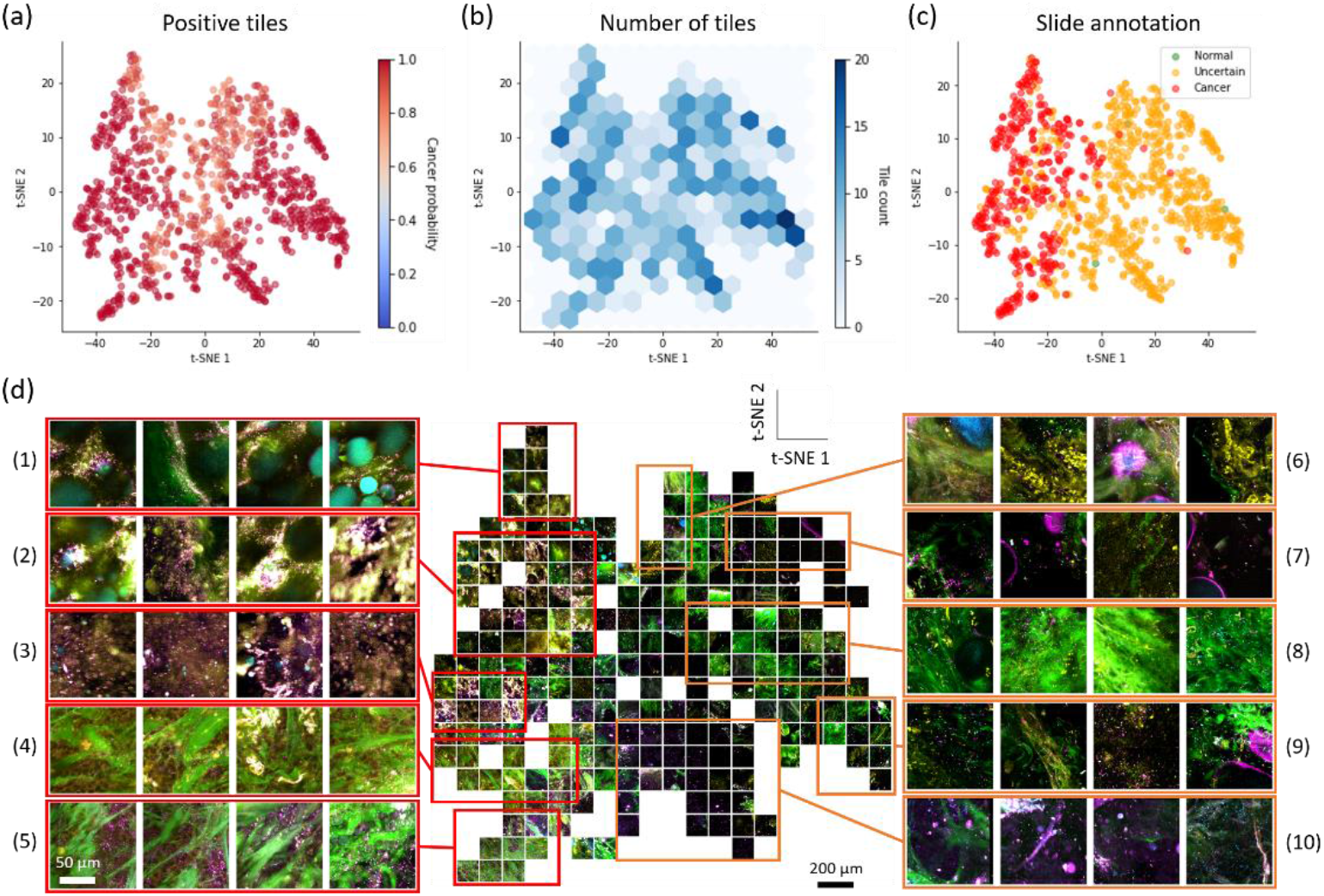
Visualization of the latent feature representations of predicted positive tiles (cancer probability > 0.5). (a) t-SNE plot of positive tiles predicted by MM-MIL. (b) A hexagonal heatmap showing the number of tiles in each region of the t-SNE plot. (c) Cancer-related patterns from “Cancer” and “Uncertain” SLAM slides exhibit both similarities (left part, with tiles from both “Cancer” and “Uncertain” slides) and differences (right part, with mainly tiles from “Uncertain” slides). (d) Positive SLAM tiles corresponding to the dots in the t-SNE plot were sampled and visualized in the same locations. Positive tiles in the red bounding boxes (1)-(5) came from “Cancer” and “Uncertain” slides, while positive tiles inside orange bounding boxes (6)-(10) came predominantly from “Uncertain” slides.

The corresponding tile images in the t-SNE plot are visualized in Fig. 6(e), with each channel of the images (i.e., SHG, THG, 2PF, 3PF) to be visualized separately in Fig. S4 in Supplement 1. It can be observed that the normal patterns presented in negative tiles (cancer probability ≤ 0.5) include collagen (green, from second harmonic generation - SHG) and adipocytes (cyan, from three-photon fluorescence - 3PF). While in positive tiles, significant optical heterogeneity is observed, implicating the existence of various cancer-related patterns. This is further demonstrated in Fig. 7, where the latent feature space of positive tiles was visualized. The apparent discrepancy of cancer-related patterns from “Cancer” and “Uncertain” slides can also be observed in Fig. 7(c), where tiles from both “Cancer” and “Uncertain” slides appear on the left half of the t-SNE plot, while positive tiles on the right mainly are from “Uncertain” slides. As shown in Fig. 7(d), predicted positive tiles show the diversity of optical characteristics of cancer-related tiles revealed by MM-MIL. From these observations, we can infer that MM-MIL is able to identify cancer-related patterns from both “Uncertain” and “Cancer” slides to differentiate positive bags from the negative ones during the training process. This property makes MM-MIL useful for optical signature discovery in uncertain areas, where cancer-related patterns may not exist in “Cancer” slides.

### 3.4 Channel-wise pixel-level model interpretation

Considering that the four detection modalities of SLAM provide distinct optical contrasts of tissue samples, learning the contribution of each SLAM channel may inspire the discovery of the cancer-related optical signatures. Here, an occlusion-based method was utilized to inform the importance of each SLAM channel for cancer prediction. In addition, saliency maps were generated, which inform the prominent structures on the pixel level. It was observed that the highly informative channels (the ones inducing significant change in prediction score after occlusion) differ among positive tiles (Fig. 8), which indicates that the importance of each channel for cancer prediction may vary within positive tiles. Among them, 2PF, THG, and SHG channels have high contribution for cancer tile predictions in “Cancer” slides [Fig. 8(a-f)]. While for the positive tiles in “Uncertain” slides, the 3PF channel has predominant contribution to the predictions [Fig. 8(g-1)]. According to the saliency maps, the cyan point-like objects in the 3PF channel were determined as a highly informative feature for cancer prediction. Those cancer-related patterns were further discussed in the following sections.

**Fig. 8.**
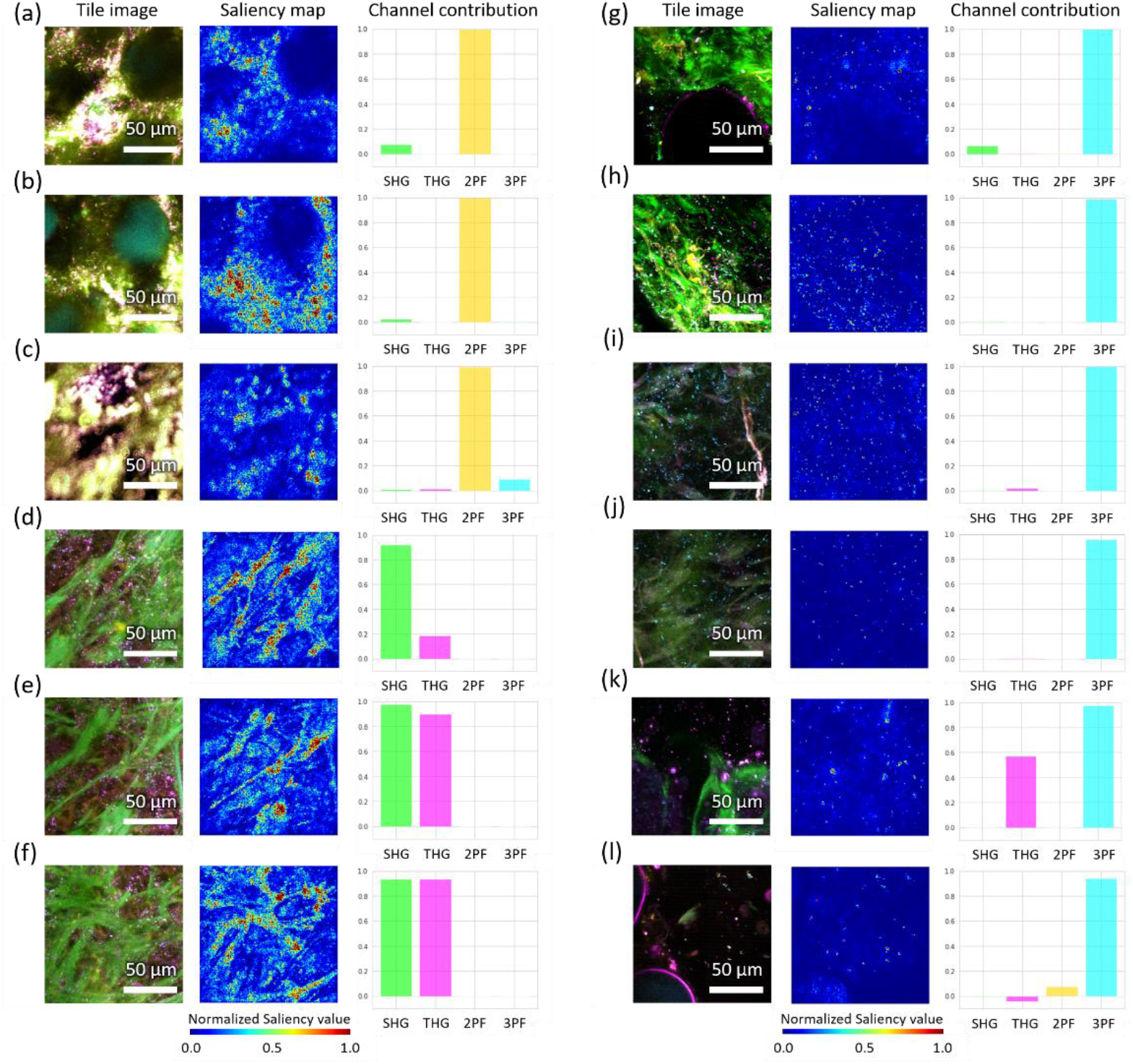
Cancer-related patterns informed by saliency maps and channel occlusion measurements of predicted positive tiles. The values in the channel occlusion bar charts represent the decrease in predicted cancer probability when a particular SLAM channel is occluded. Representative cancer tiles from “Cancer” slides are shown in (a-f). Representative positive tiles from “Uncertain” slides are shown in (g-1).

### 3.5 Cancer-related patterns informed by MM-MIL

Based on the forementioned localization predictions and model interpretation methods, several cancer-related patterns were revealed by MM-MIL, including well-known cancer biomarkers and non-obvious cancer-related patterns.

#### 3.5.1 Well-known cancer biomarkers

Dense clusters of tumor cells are visible in the positive tiles from “Cancer” slides [Fig. 7(d), rows (4) and (5)]. Those breast cancer tumor cells are reported to have high THG signal intensity (magenta colored) in SLAM images [34], which match with our findings that the THG channel has a significant contribution to cancer prediction as shown in Fig. 8(d-f).

Additionally, collagen fibers (visible in SHG channel) are highlighted in positive tiles [Fig. 8(d-f)]. In breast cancer, stromal collagen fibers participate in the migration of metastatic tumor cells which may promote tumorigenesis [35]. Based on the morphology and their interactions with tumor cells, collagen fiber patterns in breast cancer can be categorized into different types of tumor-associated collagen signatures (TACS) [36, 37]. Using the saliency maps and channel occlusion results, collagen fibers that co-occur with tumor cells were determined as prominent structures for cancer diagnosis by MM-MIL [Fig. 8(d-f)]. This specific pattern matches with the characteristics of TACS-6, which is defined by the disordered alignment of collagen fibers that enables multidirectional tumor cell migration. According to pathological reports, such collagen fiber patterns appeared in SLAM images from subjects who were diagnosed with invasive ductal carcinoma. This aligns the previous observation that TACS-6 appears at the invasive stage of tumor development [37]. Constrained by the FOV of tiles at all magnification levels (i.e., 128 μm × 128 μm for high, 256 μm × 256 μm for medium, and 512 μm × 512 μm for low-magnification level), other large-scale TACS signatures (mainly defined on the millimeter scale) could not be observed in MM-MIL predictions.

#### 3.5.2 Non-obvious cancer-related patterns

The non-obvious cancer-related patterns, which mainly come from “Uncertain” slides, are visualized in [Fig. 7(d), rows (6-10)]. Among those patterns, we found that the cyan point-like objects in the 3PF channel were repeatedly highlighted by saliency maps [Fig. 8(g-l)]. Occluding the 3PF channel leads to a significant drop in the prediction score. Such pattern can also be observed as clusters among positive tiles in the latent feature space (Fig. S4).

Previous studies showed that these cyan dots are NAD(P)H-rich extracellular vesicles (EVs), which appear as diffraction-limited punctuated pixels in the SLAM 3PF channel [38, 39]. EVs play an important role in intercellular communication between cancer cells and the tumor microenvironment [38]. They contribute to cancer growth and metastasis with multiple functionalities, including the suppression of immune response, the recruitment of stromal cells, and the determination of organotrophic metastasis [40, 41]. The high concentration of NAD(P)H has also been reported in cancer cells, which is considered to be linked to the antioxidant defense mechanism [42] and Warburg effect [43]. In addition, EVs from breast cancer cells are reported to have significantly higher NAD(P)H concentration compared with EVs from nontumorigenic cells [39]. It is also reported that EVs outside the visible tumor area (as far as 5 cm away from the tumor boundary) show even higher NAD(P)H concentration compared to EVs within the tumor, demonstrating the far-reaching impact of EVs in carcinogenesis [44, 45]. Those reported results support the findings by MM-MIL that NAD(P)H-rich EVs (or cyan dots in the 3PF channel) are associated with breast cancer, and that such pattern can be more easily observed in peri-tumoral regions (mainly represented by “Uncertain” slides) than tumor areas (shown in the “Cancer” slides).

In addition to forementioned patterns, we also observed some suspicious cancer-related patterns that are difficult to attribute to known biological processes or mechanisms, including the FAD-rich punctuated patterns surrounding adipocytes [Fig. 8(a-c)]. Further studies are needed to verify the correlation between these unveiled optical signatures and human breast cancer.

## 4. Discussion

In this study, we developed a weakly-supervised deep learning framework (MM-MIL) for the discovery of human breast cancer-related optical signatures based on label-free virtual histopathology (SLAM microscopy) which contains rich structural and functional information in tissue samples. The task of this optical signature discovery has three main characteristics: (1) inexact supervision, (2) incomplete supervision, and (3) positive instances in labeled and unlabeled data that may come from separate inherent distributions. MM-MIL combines the inexact and incomplete supervision into one MIL task via the Mix-and-Match bagging policy, which ensures the explicitness of bag-level labels. The established framework detected cancer-related patterns in both primary tumor areas and surrounding peri-tumoral regions. Based on model interpretation, a variety of human breast cancer-related patterns were revealed from SLAM virtual histopathology slides, including well-known cancer signatures as well as non-obvious patterns. Among them, the presence of NAD(P)H-rich EVs support our previous suggestion that breast-cancer-associated EVs could have extensive impact on carcinogenesis, which would be associated with field cancerization [39, 46]. The optical signatures found by MM-MIL inspire new hypotheses for cancer biomarker discovery and translational clinical applications, and merit further investigation.

MM-MIL is capable of learning cancer-related patterns from both labeled and unlabeled data. Instead of taking shortcuts by making positive predictions only on “Cancer” slides, MMMIL tracked cancer-related patterns in both “Cancer” and “Uncertain” slides. This can be explained by the fact that during the early stage of the training process, the selection of discriminative tiles for model weights updates is mainly stochastic. Since slide-level labels are hidden during training, it is unlikely for the model to pick tiles only from “Cancer” slides under the circumstances that “Uncertain” slides also contain cancer-related patterns. In general, the goal of MM-MIL is to reveal patterns that are not observable from “Normal” slides, regardless of the slide-level labels (i.e., “Cancer” or “Uncertain” slides).

Given the uniqueness of the multi-modal images (i.e., each channel of SLAM provides distinct structural and functional information about the tissue), knowing the contribution of each channel and the salient structures is beneficial for the understanding of model behavior. Thus, two channel-wise model interpretation methods (i.e., channel occlusion and saliency maps) were leveraged. Using these techniques, several well-known cancer biomarkers as well as non-obvious cancer-related patterns were revealed in both the primary tumor areas and the surrounding peri-tumoral regions, which align with the before-mentioned assumption of potentially separate distributions of positive instances.

The MM-MIL framework has some limitations. The cancer-related patterns learned by MM-MIL are constrained by the size of the FOV of the tile images. In this study, we focus on patterns at the scale of 128 μm, 256 μm, and 512 μm. To learn larger-scale patterns (e.g., large-scale tumor-associated collagen signatures), larger tile FOVs would be recommended. Further improvements can be made to enable optical signatures discovery for multi-class scenarios (e.g., cancer prognosis) with enhanced capability in extracting correlations of patterns across regions and scales. In addition, further validation of our framework is needed to verify its generalizability on data from different microscopy systems.

## 5. Conclusion

The proposed weakly-supervised deep learning framework offers a practical solution to gain insights into multiplexed optical bioimages when explicit annotations are unavailable. With the advancements in optical imaging techniques, many crucial biological questions can be answered based on the rich structural and functional information revealed from samples. It is widely believed that DL is proficient at extracting knowledge from large volume of data. Nevertheless, strong supervision is often difficult to obtain for optical signature discovery tasks. This study demonstrates the capability of the proposed framework in recognizing patterns and extracting correlations between optical features with human breast cancer based on inexact and incomplete supervision, which could be extended to various types of biological conditions and imaging modalities.

## Acknowledgments

The authors thank all the human subjects who consented to having their tissue be part of our imaging studies, as well as all the staff at the Carle Research Office for assisting in the coordination and consenting of subjects. We thank Sixian You, Eric J. Chaney for their technical support. We thank all the team members from the Biophotonics Imaging Laboratory for their suggestions and support. Additional information can be found at: http://biophotonics.illinois.edu.

## Data availability

The SLAM human breast cancer datasets that support the findings of this study are available from the corresponding author (S.A.B.) upon request and through collaborative investigations.

## Supplemental document

See Supplement 1 for supporting content.

