## Supplement 1 for "Weakly supervised identification of microscopic human breast cancer-related optical signatures from normal-appearing breast tissue"

### 1. Supplementary notes

#### *1.1 EM-based Multiple Instance Learning*

In this study, we modeled the cancer signature discovery task as a Multiple Instance Learning (MIL) problem. Collections of SLAM slides grouped by the Mix-and-Match bagging policy were treated as bags, whereas the tile images extracted from SLAM slides were treated as instances. We denote $B = \{B_1, B_2, \ldots, B_N\}$ as the dataset containing $N$ bags. Each bag $B_i = \{x_{i,1}, x_{i,2}, \ldots, x_{i,N_i}\}$ has $N_i$ instances. The corresponding ground-truth labels are $Y = \{Y_1, Y_2, \ldots, Y_N\}$ , where all the instances in each bag share the same labels $Y_i = \{y_{i,1}, y_{i,2}, \ldots, y_{i,N_i}\}, y_{i,j} = y_i$. Here we introduce hidden variables $H = \{H_1, H_2, \ldots, H_N\}$, $H_i = \{H_{i,1}, H_{i,2}, \ldots, H_{i,N_i}\}$, where $H_{i,j}$ indicates whether instance $x_{i,j}$ is discriminative for label $y_i$. We assume that the bags $B$ are independent and identically distributed, and that instance $x_{i,j}$ depends only on $H_{i,j}$ while being independent with each other given $H_{i,j}$.Thus we can write the generative model that generates $B, Y$ and $H$ as:

$$P(B, Y, H) = \prod_{i=1}^{N} \prod_{j=1}^{N_i} (P(x_{i,j}, y_i | H_{i,j}) P(H_i)). \tag{S1}$$

The data likelihood $P(B, Y, H)$ can then be maximized using the EM algorithm. The optimization process consists of E-steps and M-steps. For the initial E-step, $H_{i,j}$ is randomly set to 1 for instances, meaning that the discriminative instances in positive and negative bags are randomly chosen. During the M-steps, the parameter $\theta$ of the instance-level classifier ($f_{ins}$) is updated to maximize the data likelihood based on the set of discriminative instances ($T$):

$$\begin{aligned} \theta &\leftarrow \underset{\theta}{\operatorname{argmax}}\, P(B, Y | H; \theta) \\ &= \underset{\theta}{\operatorname{argmax}} \prod_{x_{i,j} \in T} P(x_{i,j}, y_i | \theta) \\ &= \underset{\theta}{\operatorname{argmax}} \prod_{x_{i,j} \in T} P(y_i | x_{i,j}; \theta) P(x_{i,j} | \theta) \\ &= \underset{\theta}{\operatorname{argmax}} \prod_{x_{i,j} \in T} P(y_i | x_{i,j}; \theta), \end{aligned} \tag{S2}$$

which describes a discriminative model. Here we assume that non-discriminative instances are generated from a uniform generative model, and that $x_{i,j}$ is from a uniform distribution. In the following E-steps, the hidden variables $H$ are estimated by $f_{ins}$. Based on the standard Multi-instance assumption [1], the instance with the maximum $P(H_{i,j} | x_{i,j})$ is discriminative in each bag. In other words, the instance-level classifier aims to differentiate the most-likely positive instances in positive MIL bags from the least-likely negative instances in negative MIL bags.

#### *1.2 Instance-level classifier optimization*

The optimization of the instance-level classifier ( $f_{ins}$ ) in MM-MIL is described in Algorithm S1. The sets of training and validation bags are denoted as $B_{train}$ and $B_{val}$. The bag-level ground truth labels are denoted as $Y_{train}$ and $Y_{val}$ for training and validation sets respectively, whereas $\hat{Y}_{train}$ and $\hat{Y}_{val}$ represent the bag-level predictions. $K$ (number of instances to be selected from each bag) and $N_{epoch}$ (number of training epochs) are user-defined parameters. $\mathcal{L}_{train}$ is the training loss (i.e., cross-entropy loss).

---

**Algorithm S1**: Optimization of the instance-level classifier

---

**input:** $B_{train}$, $Y_{train}$, $B_{val}$, $Y_{val}$, $f_{ins}$, $K$, $N_{epoch}$

**output:** trained model ($f_{ins}$), validation results ($records$)

$epoch$ = 1

$records$ = {}

**while** $epoch \leq N_{epoch}$ **do**

/* inference step */

$\hat{Y}_{train}$= model_inference ($B_{train}$, $f_{ins}$)

$T_{train}$ = select_instances($B_{train}$, $Y_{train}$, $\hat{Y}_{train}$, $K$)

/* training step */

$f_{ins}$, $\mathcal{L}_{train}$ = train_model($f_{ins}$, $T_{train}$)

/* validation */

$\hat{Y}_{val}$ = model_inference($B_{val}$, $f_{ins}$)

$metrics$ = prediction_evaluation($Y_{val}$, $\hat{Y}_{val}$)

$records$ = {$records, (epoch, \mathcal{L}_{train}, metrics)$}

/* early stopping */

$flag$ = determine_early_stopping($records$)

**if** $flag$ = $true$ **then**

**return** $f_{ins}$, $records$

**end if**

$epoch$ = $epoch$ + 1

**end while**

**return** $f_{ins}$, $records$

#### 1.3 Visualization of latent feature space

To visualize the latent feature space of the model, we randomly sampled 1,500 tile images from the test set, in addition to all predicted positive tiles (probability > 0.5) by the MM-MIL model that was trained on the high-magnification level. The latent feature representations of those tiles were extracted from the output of the adaptive average pooling layer of the ResNet34 model. The feature space was visualized in two dimensions using t-distributed stochastic neighbor embedding (t-SNE) [2]. SLAM tiles (converted to RGB images) corresponding to the points in the t-SNE plot were sampled and visualized for model introspection.

#### 1.4 Occlusion-based channel contribution measurement

To investigate the relevance of each individual SLAM channel for cancer prediction, we occluded each channel of the positive tiles and measured the decrease of prediction score, which can be seen as an indicator of channel importance regarding to the prediction task. The more the prediction score dropped, the more relevant the channel would be to cancer prediction. To eliminate the effect of out-of-distribution (OOD) test examples, the channel occlusion was achieved by replacing the targeted channel with Gaussian noise that imitates uninformative signals in that channel. The mean and variance of the Gaussian noise were measured based on manually annotated background areas in 460 predicted positive tiles. When tested on trained MM-MIL models, the predicted cancer probabilities of the Gaussian noise input were close to 0 ($7.92\times10^{-7}\pm3.41\times10^{-8}$), compared to 0.72±0.04 when using an all-zero input (mainly due to OOD).

#### 1.5 Saliency maps for pixel-level model interpretation

To gain insights into the salient structures on the pixel-level, saliency maps were generated using Integrated Gradients (IG) [3]. The baseline input of IG was Gaussian noise (same as the informative signal used in channel occlusion). To reduce the visual noise in the attribution maps, SmoothGrad (SG) [4] was implemented, with the number of randomly generated examples set to be 5, and the standard deviation of SG Gaussian noise to be 0.02.

### 2. Supplementary Figures

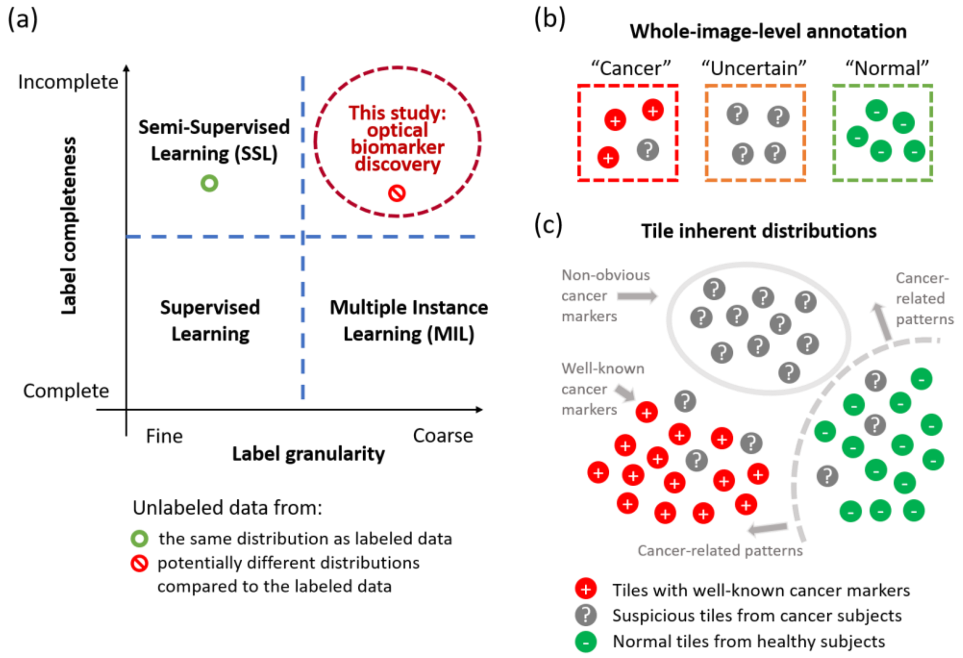


**Fig. S1**. Description of the task of optical signature discovery in the study. (a) This study focuses on the identification of human breast cancer-related optical signatures by leveraging inexact and incomplete supervision. (b) Whole-image-level annotations were generated for SLAM virtual histology slide images. (c) A diagram showing the potential inherent distributions of SLAM tile images. We define the cancer-related patterns as the optical characteristics that appear only in tissues from cancer subjects, while being unobservable in normal tissues from healthy subjects. The potentially new optical biomarkers (non-obvious cancer-related optical signatures) may emerge from different distributions compared to well-known cancer markers.

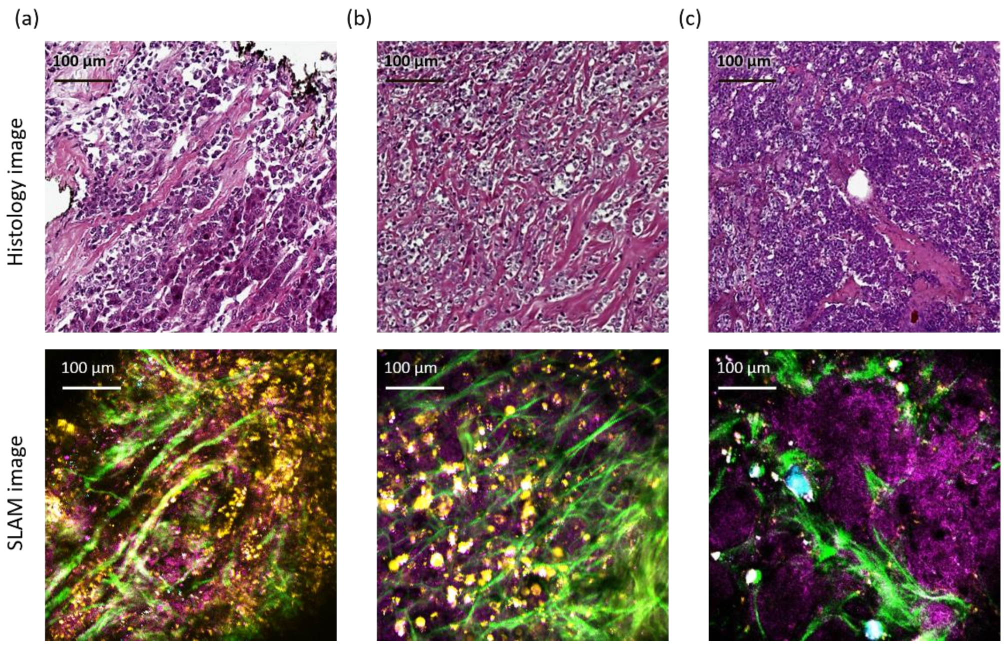


**Fig. S2.** SLAM virtual histopathology images and the corresponding histology of different types of breast cancer. (a) Invasive ductal carcinoma (IDC). (b) Invasive lobular carcinoma (ILC). (c) Invasive micro-papillary carcinoma (ImPC).

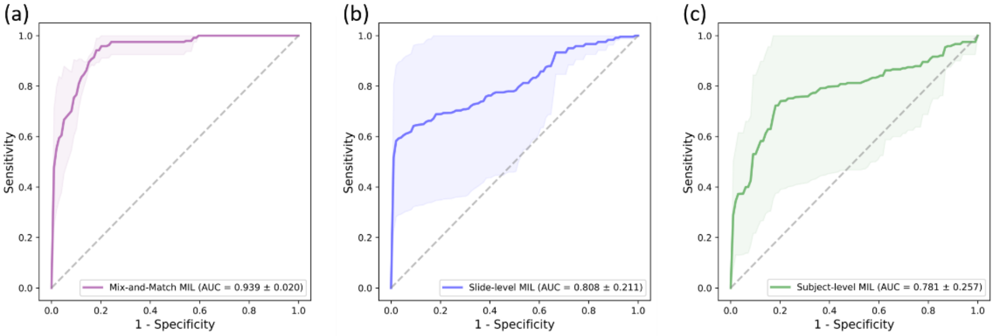


**Fig. S3**. ROC curves of MIL models with different bagging policies evaluated on slides with certain labels (“Cancer” and “Normal”). The solid lines indicate the mean curve over five models trained under the same configuration, and the shaded regions indicate ± standard deviation. (a) MIL models trained on bags generated via the Mix-and-Match bagging policy. (b) Each SLAM slide image was treated as an individual bag. Slides from cancer subjects were treated as positive bags, otherwise they were treated as negative bags. (c) Slides from each subject were grouped into a bag (positive for cancer subjects, and negative for normal subjects).

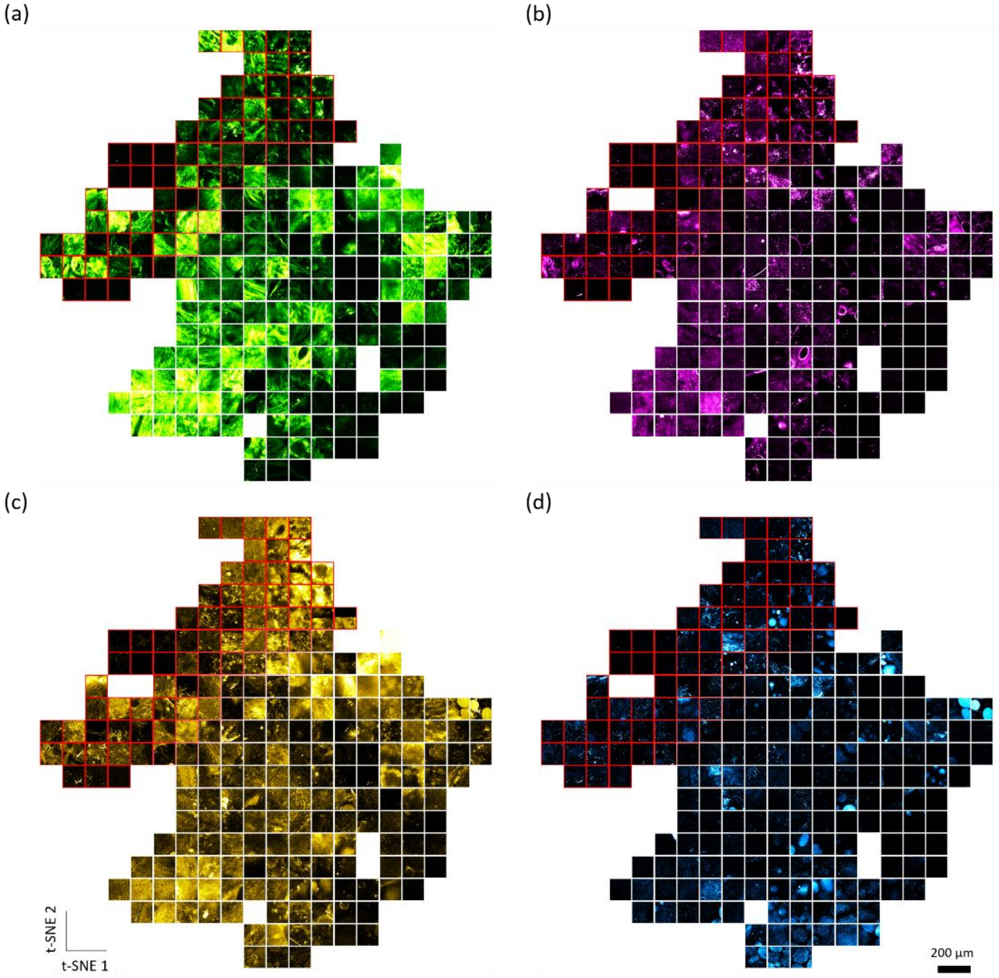


**Fig. S4**. t-SNE visualization of the latent feature space of a trained MM-MIL model. Each channel of the SLAM images is visualized separately: (a) SHG channel, (b) THG channel, (c) 2PF channel, and (d) 3PF channel.

### 3. Supplementary Tables

**Table S1. Demographic information on human subjects, pathological diagnoses, and SLAM imaging sights. This study included 36 breast cancer subjects (S-001 to S-036) and 12 healthy cancer-free subjects (S-037 to S-048). IDC, invasive ductal carcinoma; DCIS, ductal carcinoma *in situ*; ILC, invasive lobular carcinoma; LCIS, lobular carcinoma *in situ*; IPC, invasive papillary carcinoma. For subject S-001 and S-002, who had IDC before the neoadjuvant therapy prior to surgery, primary tumor was not found in their tissue samples. Their SLAM slides were labeled as "Uncertain".**

| Subject ID | Age | Pathologic Stage | Type of surgery | Type of cancer | # SLAM slides |
|---|---|---|---|---|---|
| S-001 | 48 | 0 | Bilateral breast mastectomy | No primary tumor found | 36 |
| S-002 | N/A | 0 | Bilateral breast mastectomy | No primary tumor found; fibroadenoma | 20 |
| S-003 | 51 | 0 | Left breast lumpectomy | DCIS | 16 |
| S-004 | 82 | I | Bilateral breast mastectomy | IPC, DCIS | 28 |
| S-005 | 65 | I | Right breast mastectomy | IDC, DCIS | 24 |
| S-006 | 76 | I | Right breast lumpectomy | IDC, DCIS | 24 |
| S-007 | 47 | I | Bilateral breast mastectomy | IDC, LCIS, DCIS | 8 |
| S-008 | 68 | I | Right breast lumpectomy | IDC, DCIS | 8 |
| S-009 | 52 | I | Right breast mastectomy | IDC, LCIS, DCIS | 16 |
| S-010 | 38 | I | Right breast mastectomy | IDC, DCIS | 4 |
| S-011 | 71 | I | Right breast mastectomy | IDC, DCIS | 4 |
| S-012 | 57 | I | Right breast lumpectomy | IDC, DCIS | 8 |
| S-013 | 64 | I | Right breast lumpectomy | IDC | 16 |
| S-014 | 52 | I | Left breast mastectomy | IDC, DCIS | 4 |
| S-015 | 61 | I | Left breast lumpectomy | IDC, LCIS | 12 |
| S-016 | 35 | I | Right breast mastectomy | IDC, DCIS | 8 |
| S-017 | 53 | I | Right breast mastectomy | IDC, DCIS | 12 |
| S-018 | 76 | II | Right breast mastectomy | IPC, DCIS | 16 |
| S-019 | 56 | II | Bilateral breast mastectomy | IDC, DCIS | 24 |
| S-020 | 71 | II | Bilateral breast mastectomy | IDC | 20 |
| S-021 | 82 | II | Right breast mastectomy | IDC, DCIS | 32 |
| S-022 | 81 | II | Right breast mastectomy | IDC, DCIS | 12 |
| S-023 | 89 | II | Left breast mastectomy | ILC, IDC, DCIS | 4 |
| S-024 | 73 | II | Right breast lumpectomy | IDC, DCIS | 4 |
| S-025 | 71 | II | Right breast lumpectomy | IDC, DCIS | 20 |
| S-026 | 53 | II | Right breast lumpectomy | IDC, DCIS | 8 |
| S-027 | 59 | III | Left breast mastectomy | IDC | 4 |
| S-028 | 46 | III | Right breast mastectomy | IDC, DCIS | 28 |
| S-029 | 77 | III | Left breast mastectomy | ILC, LCIS, DCIS | 16 |
| S-030 | 64 | III | Left breast mastectomy | IDC, DCIS | 16 |

| | | | | | |
|---|---|---|---|---|---|
| S-031 | 63 | N/A | N/A | N/A | 8 |
| S-032 | 60 | N/A | Left breast mastectomy | ILC, DCIS | 20 |
| S-033 | 57 | N/A | Prophylactic mastectomy | LCIS, lobular hyperplasia | 12 |
| S-034 | 43 | Benign | Right breast lumpectomy | Benign | 8 |
| S-035 | 56 | Benign | Right breast lumpectomy | LCIS | 8 |
| S-036 | 62 | Benign | Left breast lumpectomy | Benign | 16 |
| S-037 | 19 | Normal | Breast reduction surgery | — | 8 |
| S-038 | 70 | Normal | Breast reduction surgery | — | 28 |
| S-039 | 40 | Normal | Breast reduction surgery | — | 36 |
| S-040 | 56 | Normal | Breast reduction surgery | — | 28 |
| S-041 | 50 | Normal | Breast reduction surgery | — | 24 |
| S-042 | 41 | Normal | Breast reduction surgery | — | 8 |
| S-043 | 45 | Normal | Breast reduction surgery | — | 20 |
| S-044 | N/A | Normal | N/A | — | 16 |
| S-045 | 36 | Normal | Prophylactic bilateral mastectomy | — | 8 |
| S-046 | 18 | Normal | Breast reduction surgery | — | 8 |
| S-047 | 22 | Normal | Breast reduction surgery | — | 8 |
| S-048 | 18 | Normal | Bilateral mastectomy | — | 8 |

**Table S2. Details of bags and tiles under three magnification settings.**

| | | | | |
|---|---|---|---|---|
| Tile FOV: 256×256 pixels; overlap ratio: 0% | | | | |
| Set | # Bags | # Positive bags | # Instances | # Positive instances |
| Training | 1,098 | 354 | 98,490 | 72,300 |
| Validation | 186 | 42 | 18,582 | 12,960 |
| Test | 456 | 192 | 44,316 | 32,424 |
| Tile FOV: 512×512 pixels; overlap ratio: 50% | | | | |
| Set | # Bags | # Positive bags | # Instances | # Positive instances |
| Training | 1,098 | 354 | 85,588 | 62,196 |
| Validation | 186 | 42 | 15,582 | 10,805 |
| Test | 456 | 192 | 37,578 | 27,300 |
| Tile FOV: 1024×1024 pixels; overlap ratio: 80% | | | | |
| Set | # Bags | # Positive bags | # Instances | # Positive instances |
| Training | 1,098 | 354 | 95,136 | 70,043 |
| Validation | 186 | 42 | 15,978 | 10,944 |
| Test | 456 | 192 | 40,626 | 29,226 |

**Table S3. List of parameters used in our experiments across three magnification levels.**

| Parameters | Magnification levels | | |
|---|---|---|---|
| | High | Medium | Low |
| Tile FOV (pixels) | 256×256 | 512×512 | 1024×1024 |
| Tile size (pixels) | 256×256 | 256×256 | 256×256 |
| Tile overlap ratio | 0% | 50% | 80% |
| Epoch | 200 | 200 | 200 |
| Batch size | 600 | 600 | 600 |
| Learning rate | $1.5\times10^{-4}$ | $1.5\times10^{-4}$ | $1.5\times10^{-4}$ |
| Optimizer | Adam<br>$\beta_1$, $\beta_2$ = (0.9, 0.999)<br>weight_decay = $1\times10^{-5}$ | Adam<br>$\beta_1$, $\beta_2$ = (0.9, 0.999)<br>weight_decay = $1\times10^{-5}$ | Adam<br>$\beta_1$, $\beta_2$ = (0.9, 0.999)<br>weight_decay = $1\times10^{-5}$ |
| Number of top-K instances ($K$) | 15 | 15 | 30 |
| Backbone of model $f_{ins}$ | ResNet34 | ResNet34 | ResNet34 |
| Positive class weight ($w_1$) | 0.66 | 0.65 | 0.66 |
| Negative class weight ($w_0$) | 0.34 | 0.35 | 0.34 |
| Model Selection | Highest validation accuracy | Highest validation accuracy | Highest validation accuracy |
| Validation frequency | Every 5 epochs | Every 5 epochs | Every 5 epochs |
